# Metformin disrupts *Danio rerio* metabolism at environmentally relevant concentrations: A generational study

**DOI:** 10.1101/2022.04.05.487223

**Authors:** Susana Barros, Marta Ribeiro, Ana M. Coimbra, Marlene Pinheiro, Hugo Morais, Nélson Alves, Rosa Montes, Rosario Rodil, José Benito Quintana, Miguel. M. Santos, Teresa Neuparth

**Affiliations:** CIMAR/CIIMAR—Interdisciplinary Centre of Marine and Environmental Research, Endocrine Disruptors and Emerging Contaminants Group, University of Porto, Avenida General Norton de Matos, S/N, 4450-208 Matosinhos, Portugal; CITAB - Centre for the Research and Technology of Agro-Environmental and Biological Sciences, University of Trás-os-Montes and Alto Douro (UTAD), Quinta de Prados, Pavilhão 2, 5000-801 Vila Real, Portugal; Inov4Agro –Institute for Innovation, Capacity Building and Sustainability of Agri-food Production, Portugal; Department of Analytical Chemistry, Nutrition and Food Sciences, IAQBUS - Institute of Research on Chemical and Biological Analysis, Universidade de Santiago de Compostela, Constantino Candeira S/N, 15782 Santiago de Compostela, Spain; FCUP - Department of Biology, Faculty of Sciences, University of Porto (U. Porto), Rua do Campo Alegre s/n, 4169-007 Porto, Portugal

**Keywords:** Metformin, Zebrafish, Energy metabolism, Lipid content, Generational exposure, low-level exposure

## Abstract

Metformin (MET) is an anti-diabetic pharmaceutical with a large-scale consumption, which is increasingly detected in surface waters. However, current knowledge on the generational effects of MET exposure in the metabolism of non-target organisms is limited. The present study aimed at investigating the effects of MET in the model freshwater teleost *Danio rerio*, following a generational exposure (from egg up to 9 months exposure) to environmentally relevant concentrations ranging from 361 ng/L to 13 000 ng/L. Biochemical markers were used to determine cholesterol and triglycerides levels, as well as mitochondrial complex I activity in males and females zebrafish liver. mRNA transcript changes were also assessed in the liver of both sexes by means of an exploratory RNA-seq analysis and expression levels of key genes involved in the energy metabolism and lipid homeostasis, i.e. *acaca, acadm, cox5aa, idh3a, hmgcra, prkaa1*, were determined using qRT-PCR analysis. The findings here reported revealed that MET was able to significantly disrupt critical biochemical and molecular processes involved in zebrafish metabolism, such as cholesterol and fatty acid biosynthesis, the mitochondrial electron transport chain and the tricarboxylic acid cycle, concomitantly to changes on the hepatosomatic index. Non-monotonic dose response curves were frequently detected in the gene expression profile, with higher effects observed for 361 ng/L and 2 166 ng/L concentrations. Collectively, the obtained results suggest that environmentally relevant concentrations of MET are able to severely disrupt *D. rerio* metabolism, with potential impacts at the ecological level, supporting the need to update the environmental quality standard (EQS) and predicted no-effect concentration (PNEC) for MET.

**Graphical abstract:** 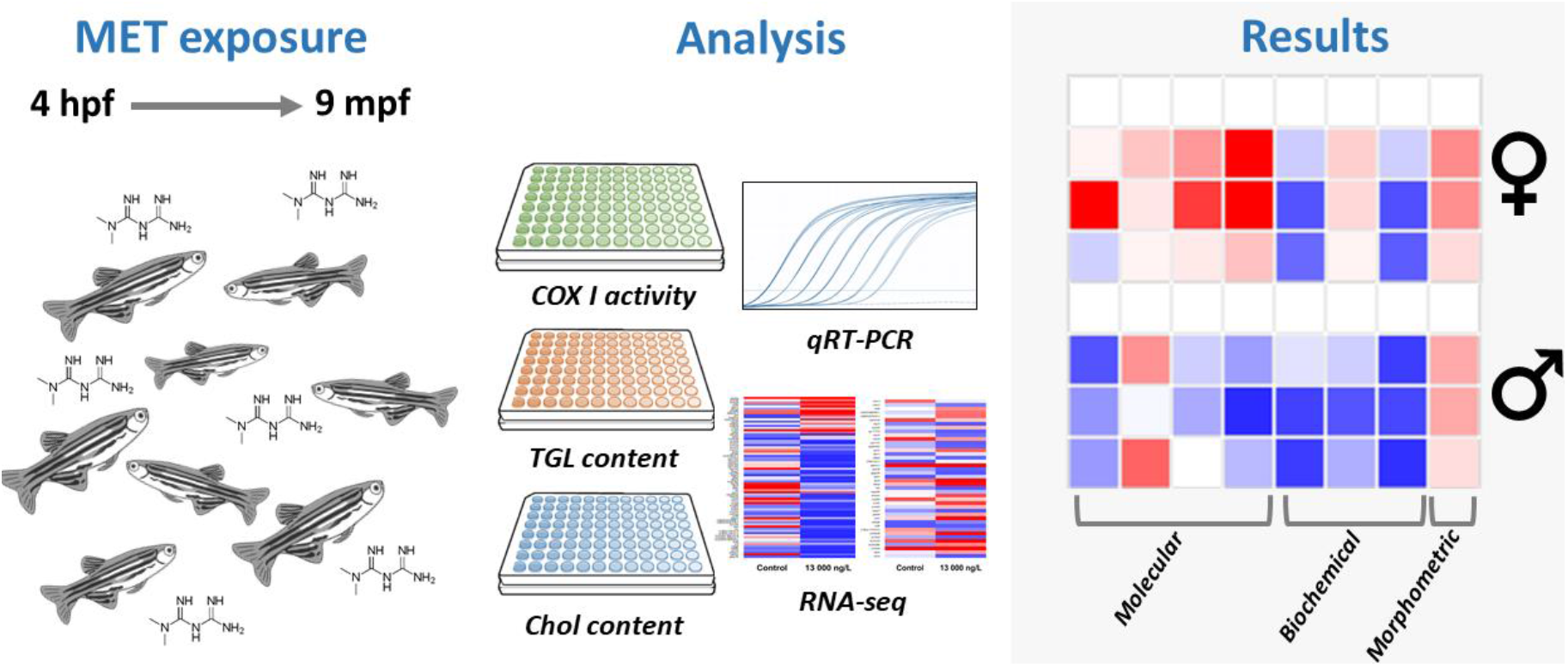

**Highlights:**

- *Danio rerio* was exposed to metformin for a full generation;
- MET affected COX I activity, as well as Chol and TGL content in zebrafish liver;
- MET altered mRNA levels of genes involved in energy metabolism and lipid content;
- Non-monotonic dose-response curves were frequently detected;
- Due to the results obtained, MET PNEC should be reviewed.

## 1. Introduction

Diabetes mellitus (DM) is one of the most prevalent chronic disease in today’s developed and developing countries, affecting around 463 million people (WHO, 2016). Currently, the predominant form is the type 2 diabetes mellitus (T2DM), representing about 90% of all diabetes cases (WHO, 2016). T2DM is generally characterized by high blood glucose content caused by insulin resistance, meaning that cells are unable to properly respond to normal levels of insulin (Adak et al., 2018).

Metformin (MET), an anti-hyperglycemic drug of the biguanides’ class, is the first-line oral therapy for T2DM patients since 2009, and one of the most prescribed pharmaceuticals worldwide (Adak et al., 2018; Thomas & Gregg, 2017). MET mode of action (MoA) in humans is not yet fully understood, however it is reported to directly inhibit the complex I of the electron transport chain, which will decrease ATP production and lead to AMP accumulation, ultimately activating the energy-sensing AMP-activated protein kinase (AMPK) (Adak et al., 2018; Sanders et al., 2007; Shurrab & Arafa, 2020). Alterations in both ATP and AMP ratios and in the AMPK pathway will lead to the suppression of hepatic gluconeogenesis, as well as the synthesis of lipids and cholesterol. Simultaneously, glucose uptake into muscle cells is improved and fatty acid ß-oxidation and glycolysis are stimulated (Figure 1) (Adak et al., 2018; Foretz et al., 2019; Shurrab & Arafa, 2020).

**Figure 1.**
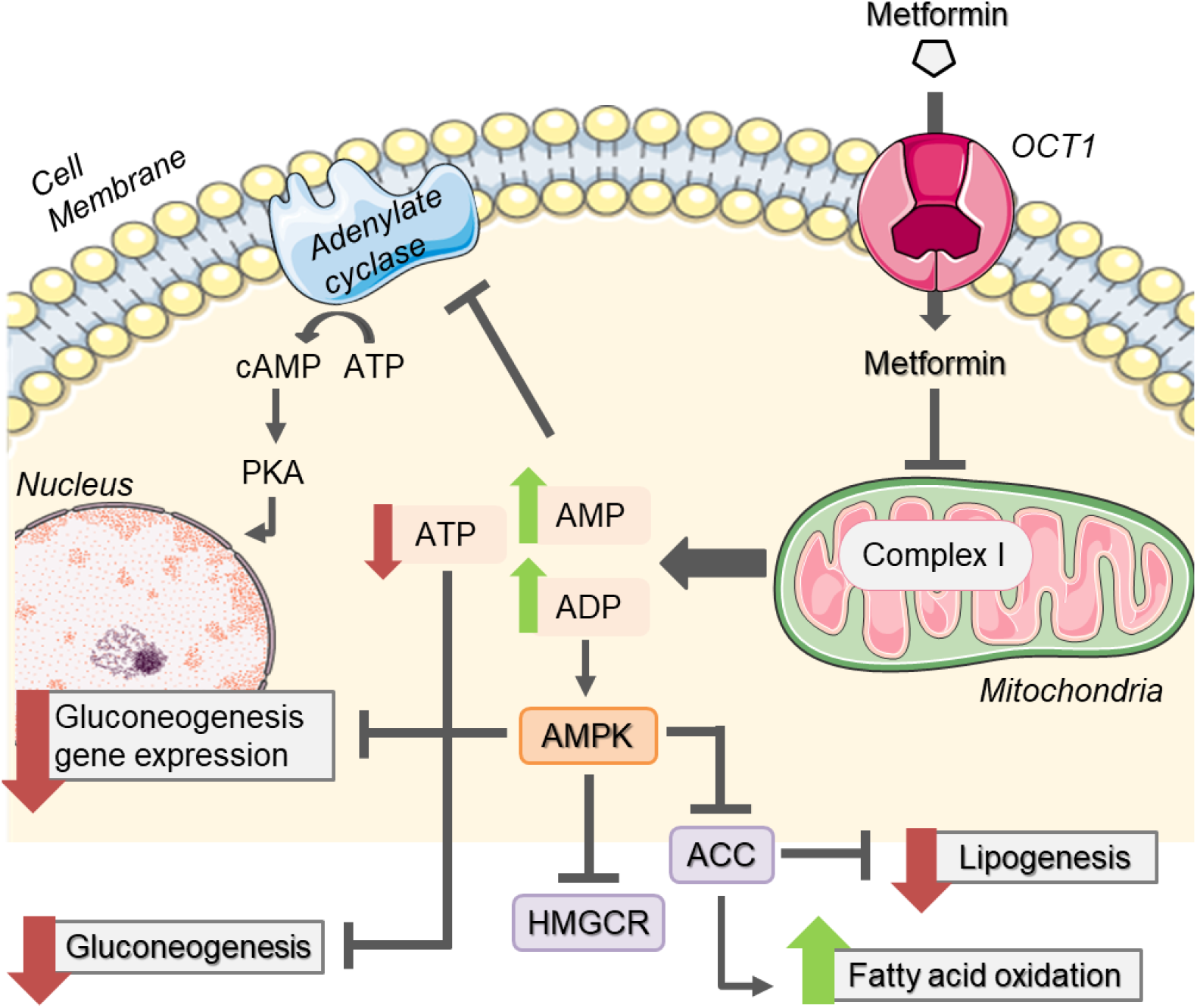
Metformin putative mode of action in humans. OCT1 - Organic cation transporter 1; AMP - adenosine monophosphate; ADP - adenosine diphosphate; ATP - adenosine triphosphate; AMPK - AMP-activated protein kinase; ACC - acetyl-CoA carboxylase; HMGCR - 3-hydroxy-3-methyl-glutaryl-coenzyme A reductase; cAMP – cyclic adenosine monophosphate; PKA - protein kinase A.

Once consumed, MET is not metabolized by the human body being excreted in its native form, mainly through urine (Bailey, 2017; Foretz et al., 2014). Due to its massive consumption worldwide, increasing levels of MET are commonly detected in wastewater treatment plants (WWTPs) influents in concentrations as high as 720 µg/L in Canada (Oliveira et al., 2015) and 325 µg/L in Portugal (de Jesus Gaffney et al., 2017). Additionally, most conventional WWTP do not eliminate completely several pharmaceuticals, including MET (Zhu et al., 2017), leading to the presence of this compound in the effluents, which in turn will be discharged into the aquatic environment (Ortiz de García et al., 2014). MET has been detected in surface waters around the world, at levels as high as 5.4 µg/L in China (Yao et al., 2018), 10.1 µg/L in Canada (de Solla et al., 2016), and 33.6 µg/L in the USA (Elliott et al., 2017). Additionally, a recent study of ours detected an average MET concentration of 852 ng/L (from 138 to 3 282 ng/L) in several rivers in the north of Portugal and Galicia (unpublished data).

Considering its high environmental concentrations in surface waters, MET has been raising concerns regarding its possible detrimental effects in non-target aquatic species. As such, MET has already been preselected in the draft of the 4^th^ watch list under the water framework directive (European Commission et al., 2022). However, the environmental quality standard (EQS) and predicted no effect concentration (PNEC) form MET has been determined at 1030 µg/L and 160µg/L in the draft of the 4^th^ watch list (European Commission et al.,2022), and some literature sources still argue that it poses no harmful effects to wildlife, at current environmental concentrations (Health and Medical Care Administration, 2021). Nevertheless, recent studies have shown that MET concentrations much lower than the EQS and PNEC are able to induce adverse effects in aquatic organisms (Lee et al., 2019; Niemuth & Klaper, 2015; Ussery et al., 2018). Although data resulting from studies performed in ecotoxicological context are still very limited, the available studies focused mainly on the effects of MET in the reproductive system of aquatic organisms, due to its potential to act as an endocrine disrupting compound in non-target organisms (Lee et al., 2019; Niemuth & Klaper, 2015; Ussery et al., 2018). Although MET mode of action is known to directly affect the energy metabolism, the number of studies addressing these effects in non-target organisms is scarce. MET has been found to cause variations in growth and weight in fish, i.e. *Salmo trutta, Oryzias latipes* and *Pimephales promelas*, exposed to high µg/L concentrations (Jacob et al., 2018; Niemuth & Klaper, 2015; Ussery et al., 2018). High glycogen levels in the liver of *Salmo trutta* were also detected after 95 days of exposure to 1 µg/L of MET (Jacob et al., 2018), as well as decreased cholesterol and triglyceride content in *Nothobranchius guntheri* (Wei et al., 2020). However, there is a knowledge gap regarding the potential effects of MET after generational exposure with concentrations similar to those found in the aquatic environment. Therefore, there is an urgent need for long-term exposure studies with environmentally relevant concentrations of MET in order to infer its potential effects in metabolic processes, as well as the implications of possible detrimental effects. The present study, focused on the effects of MET in fish metabolism, integrates a larger effort that aimed at evaluating the effects of MET on the model teleost zebrafish, after a full generation exposure to environmentally relevant concentrations (361 to 13 000 µg/L) encompassing biochemical (mitochondrial complex I activity, cholesterol and triglycerides levels) and molecular signaling pathway analysis (using qRT-PCR and RNA-seq), with apical endpoints, i.e. effects on growth and reproductive performance. Here we discuss molecular and biochemical responses associated with metabolism, while morphometric and reproductive endpoints are explored in a parallel article (Barros et al., *in prep*).

## 2. Material and Methods

### 2.1. Chemicals

1,1-Dimethylbiguanide hydrochloride (CAS: 1115-70-4), commonly known as metformin (MET), was purchased from Sigma-Aldrich (Germany). MET stock and working solutions were prepared in distilled water and stored at - 20°C.

### 2.2. Species selection

*Danio rerio*, commonly known as zebrafish, is recommended as a model species in numerous ecotoxicological studies (Fang & Miller, 2012; Kimmel et al., 1995). Due to its well described development and life-cycle, easy maintenance under laboratory and extensive available data, such as genome fully sequenced (Fang & Miller, 2012; Segner, 2009), *D. rerio* was the selected model species for this study.

### 2.3. Zebrafish generational exposure to MET

A chronic ecotoxicological bioassay was performed at “Biotério de organismos aquáticos” (BOGA) located at Interdisciplinary Centre of Marine and Environmental Research (CIIMAR) in Matosinhos, Portugal. An ethical review process carried out by CIIMAR’s animal welfare body (ORBEA, 2010/63/EU Directive) was conducted prior to the beginning of the experimental work. The bioassay was performed in compliance with the European Directive 2010/63/EU on the protection of animals used for scientific purposes, and the Portuguese ‘Decreto Lei’ 113/2013.

The experiment consisted of four treatments: a control (dechlorinated water) in triplicate and three MET concentrations in duplicate: 361 ng/L, 2166 ng/L and 13000 ng/L. The selection of these MET concentrations was based on environmental concentrations reported in literature (de Jesus Gaffney et al., 2017; Oliveira et al., 2015; Ortiz de García et al., 2014). A full generational bioassay was conducted for 9 months and started by randomly allocating 400 newly fertilized wild-type zebrafish eggs (<4 hpf), purchased from Marinnova (Portugal), in 7 L aquaria. 20 days after the beginning of the assay, 100 larvae were transferred to 30 L aquaria, where they continued development. Fish density was once again reduced from 100 to 35 individuals per aquarium once they reached 60 days, maintaining this density until the end of the bioassay. The assay was performed under a flow-through system, maintaining the water flow at 1.01 L per hour by means of two peristaltic pumps, one supplied with dechlorinated, heated and charcoal filtered tap water (ISM 444, ISMATEC), and another with stock metformin solutions (ISMATEC) (Soares et al., 2009). Aquaria were maintained with a water temperature of 28 ± 1°C, under a 14:10 h (light:dark) photoperiod, pH 7.5 ± 0.2 and a mean ammonia and nitrite concentration of 0.04 ± 0.02 mg/L and 0.03 ± 0.03, respectively. Zebrafish were fed twice a day with Tetramin (Tetra, Melle, Germany), supplemented with live brine shrimp (*Artemia* spp). The amount of food provided was adjusted according to fish development stage and size, in the same proportion to all aquaria (Barros et al., 2018; Coimbra et al., 2015; Soares et al., 2009).

### 2.4. Metformin quantification by liquid chromatography-tandem mass spectrometry

Water samples from all treatments were collected throughout the assay in order to confirm the real MET concentrations. Three sampling times were performed, (T1) at the beginning, (T2) in the middle (4 months) and (T3) at the end of the bioassay. At each time, two replicates from each treatment (bulk samples from each replicate) were collected and stored at -20°C until further analysis. MET quantification in water samples from each sampling time was determined by Liquid Chromatography - Tandem Mass Spectrometry (LC-MS/MS) with an Acquity UPLC® liquid chromatography system from waters (Milford, MA, USA). 45 μL of each sample was directly injected into an Acclaim Trinity P1 column (50 mm × 2.1 mm, 3µm particle size) (Thermo Fisher Scientific, USA) maintained at a constant temperature of 35 °C and using a flow rate of 0.2 mL/min. Mobile phases consisted of (A) ultrapure water with 2% of acetonitrile and 2 mM ammonium acetate at pH 5.5 and (B) acetonitrile with 20% of ultrapure water and 20 mM ammonium acetate at pH 5.5. The applied gradient was as follows: 0–2 min, 0% B; 2–10 min, linear gradient to 100% B; 10–15 min, 100% B and finally 15–20 min, 0% B (Montes et al., 2019) The system was interfaced to a triple quadrupole mass spectrometer (XEVO TQD®) equipped with an electrospray interface (ESI). Argon was used as collision gas and Nitrogen as a nebulizing and drying gas. MET was determined in the electrospray positive polarity and multiple-reaction monitoring (MRM) mode of acquisition. Two MRM transitions were used: m/z 130 to m/z 60 as quantification transition and m/z 130 to m/z 71 as qualification transition. Under these chromatographic conditions, metformin retention time was 8.1 min. The method assured quantification limits (LOQ) of 10 ng/L. Quantification was performed by matrix matched calibration using standards prepared in dechlorinated system water in the 10-200 000 ng/L range (which was checked to be linear, R^2^=0.9992). The relative standard deviation (RSD) of MET quantification was checked at 500, 5 000 and 50 000 ng/L levels, and found to be less than 10 % in all cases, revealing good replicability of the procedure.

### 2.5. Sampling

At the end of the generational bioassay (9 months), all animals were euthanized with an overdose of 300 mg/L tricane methanesulfonate (MS-222), buffered with the same amount of sodium hydrogen carbonate to prevent acidification of the solution. Livers of males and females of each treatment were excised and weighted in order to determine the hepatosomatic index (HIS), after which they were individually frozen in liquid nitrogen and stored at - 80ºC until biochemical analysis or preserved in RNALater (Sigma-Aldrich) at - 80ºC for gene expression analysis. Survival, reproduction and morphometric data, i.e., weight, length, reproduction, Fulton’s condition factor and HIS are presented in a parallel study (Barros et al. *in prep*).

### 2.6. Mitocondrial complex I activity

Five livers per sex from each treatment were individually homogenized in 300 µL of ice-cold phosphate buffer (25mM KH_2_PO_4_, 5 mM MgCl_2_, pH=7.5), in a Precellys homogenizer, followed by a 3 min centrifugation at 1 000 G. The supernatant was collected and centrifuged a second time for 10 min at 10 000 G, in order to obtain the mitochondrial fraction. The resulting supernatant was then discharged and the pellet was re-suspended in 30 µL of phosphate buffer. The mitochondrial rich fraction was subjected to a series of five freeze thaws in order to break mitochondria’s membranes. Overall mitochondrial protein was then quantified by optical density with a ratio of absorbance at λ260/λ280 nm using a Take3™ on a microplate reader (Biotech Synergy HT) coupled with the Gen5 software (version 2.0). Samples were then diluted to a concentration of 1 µg/µL of protein in order to normalize all samples concentrations.

The mitochondrial complex I activity was determined using the MitoCheck® Complex I activity assay (Cayman, USA), following the manufacturer protocol. COX I activity is proportional to NADH oxidation, and as such, NADH decrease was measured by reading the absorbance at 340 nm in a microplate reader (Biotech Synergy HT) coupled with Gen5 software (version 2.0). COX I activity was calculated as percentage of the control group activity, by taking into account the slope of the linear portion of each curve, using the following formula: Complex I activity (%) = (rate of target sample / rate of control)×100.

### 2.7. Lipid extraction and Cholesterol and Triglycerides quantification

Lipids were extracted from adult male and female liver samples (n=6-8), using an extraction protocol with a low toxicity solvent (Barros et al., 2018; Schwartz & Wolins, 2007). Liver tissues were individually homogenized, using a Precellys 24 homogenizer, with 10 nM PBS buffer pH 7.4, containing 10 mM EDTA (10 mg of tissue per 1 mL buffer) and two ceramic beads per tube. The homogenate of each sample was transferred (500 µL), in duplicate, to test glass vials containing 5 mL of isopropanol/hexane solution (4:1 proportion). All samples were vortexed and incubated at room temperature in the dark, with constant shaking (170/180 rpm), for 1 hour. To prevent lipid peroxidation, samples were always passed through nitrogen current before closing vials. Then, samples were washed with 2 mL of petroleum ether/hexane (1:1) solution. Vials were vortexed and put in the dark for a 10 minutes’ incubation, at room temperature. 2.5 mL of Milli-Q H2O was added to each vial, then vortexed and incubated at room temperature in the dark, with constant shaking for 20 min. The organic and inorganic phases were then separated by centrifugation at 1000 × G, for 10 min. The upper phase, which contains the lipids, was collected into new vials and evaporated to dryness under nitrogen current. At the end, dried extracts were stored at - 20°C, until cholesterol and triglycerides quantification.

Before quantification, dry extracts were re-suspended in 100 µL of isopropanol and sonicated in an ultra-sound bath (Bandelin Sonorex RK100H), for 15 minutes, at room temperature. Quantification of both parameters was performed through enzymatically colorimetric assays using Infinity Cholesterol Liquid Stable reagent and Infinity Triglycerides Liquid Stable Reagent, (Thermo scientiphic, Biognóstica, Portugal) and following the manufacturer’s protocol. Absorbance was determined at 490nm using a microplate reader (Biotech Synergy HT) coupled with Gen5 software (version 2.0). In every run, a standard curve was performed for optimal quantification and all samples were measured in duplicates. Cholesterol and triolein standards were prepared by 6 serial dilutions (from 5 to 0.156251 mg/mL for cholesterol and 2 to 0.0625 mg/mL for triolein).

### 2.8. Gene expression

#### 2.8.1. RNA isolation

Samples of both male and female livers from all the different treatments (N=6-8) were individually used to isolate total RNA via the Illustra RNAspin Mini RNA Isolation kit (GE Healthcare), according to the manufacturer’s standard protocol. RNA quantification was performed by the measurement of optical density with a Take3™ on a microplate reader (Biotech Synergy HT) coupled with the Gen5 software (version 2.0). RNA quality was verified by electrophoresis in 1.5% agarose gel and through the measurement of the ratio of absorbance at λ260/λ280 nm. All isolated RNA samples were stored at - 80°C until further use.

#### 2.8.2. RNA-seq

Pools composed of 5 liver RNA samples extracted from each sex of control and 13 000 ng/L treatments, after a 9-month MET exposure, were subjected to RNA-seq and bioinformatics analysis, obtained commercially at Novogene (United Kingdom). Due to the exploratory nature of this analysis, the 13000 ng/L MET treatment was selected as this treatment has been shown to more severely affect ecological parameters in adult zebrafish (parallel study focused on growth and reproduction - Barros et al., *in prep*). Briefly, one RNA pool of 5 livers from each treatment and sex were independently subjected to sequencing (Illumina Novaseq 6000 paired-end (2×150)). Sequenced data passed then to a bioinformatics analysis in order to determine differentially expressed genes. Briefly, raw data obtained from sequencing were subjected to a quality control analysis in order to access error rate and GC content and data were then filtered in order to remove low quality reads. Once quality control was finished, the resulting reads were mapped against the *D. rerio* reference genome (https://www.ncbi.nlm.nih.gov/assembly/GCF_000002035.6), followed by gene expression quantification. Lastly, a differential expression analysis was performed using EdgeR (v3.24.3), with analysis parameters set to: false discovery rate (FDR) p-value (padj) < 0.005 and a Log_2_(FoldChange) ≥ 1.5. Hierarchical clustering was performed to cluster FPKM (expected number of Fragments Per Kilobase of transcript sequence per Millions base pairs sequenced) values of genes. Data were clustered using Log_2_(FPKM+1) values.

#### 2.8.3. qRT-PCR: Gene selection and Primer Design

To further evaluate the effects of MET at molecular level, fluorescence-based quantitative real-time PCR (qRT-PCR) was used for all the MET concentrations (361, 2 166 and 13 000 ng/L) to quantify the transcription profiles of key genes involved in the putative MoA of MET and related to energy metabolism and lipid homeostasis (Figure 2). The protein kinase AMP-activated alpha 1 catalytic subunit (*prkaa1*) was one of the first genes selected since AMPK is thought to be activated in response to changes in the ADP:ATP and AMP:ATP ratios. The reduction of ATP levels led to the selection of two genes involved in the cellular respiration. *idh3a*, a gene encoding isocitrate dehydrogenase (IDH) which is a rate-limiting enzyme in the tricarboxylic acid cycle, and *cox5aa* (cytochrome c oxidase subunit 5Aa), which is part of the complex IV of the electron transport chain and plays a key role in the aerobic mitochondrial energy metabolism (Arnold, 2012; Barros et al., 2020; Kadenbach et al., 2000). Once AMPK is activated, catabolic processes are stimulated (e.g. fatty acid ß-oxidation) and anabolic pathways are suppressed (e.g. lipogenesis, gluconeogenesis and fatty acid synthesis). The fatty acids ß-oxidation can be carried out by the medium-chain acyl-CoA dehydrogenase (*acadm*) (Barros et al., 2020). The gene encoding acetyl-CoA carboxylase alpha (*acaca*) was selected, since the cytosolic enzyme ACC catalyzes the carboxylation of acetyl-CoA to form malonyl-CoA, an essential substrate in the regulation of fatty acids synthesis (Alves-Bezerra & Cohen, 2017; Harwood, 2005). The 3-hydroxy-3-methylglutaryl-CoA reductase protein (HMGCR) is a limiting enzyme in the biosynthesis of cholesterol, and since it was one of the biochemical parameters analyzed, we selected the gene *hmgcra* (Al-Habsi et al., 2016).

**Figure 2.**
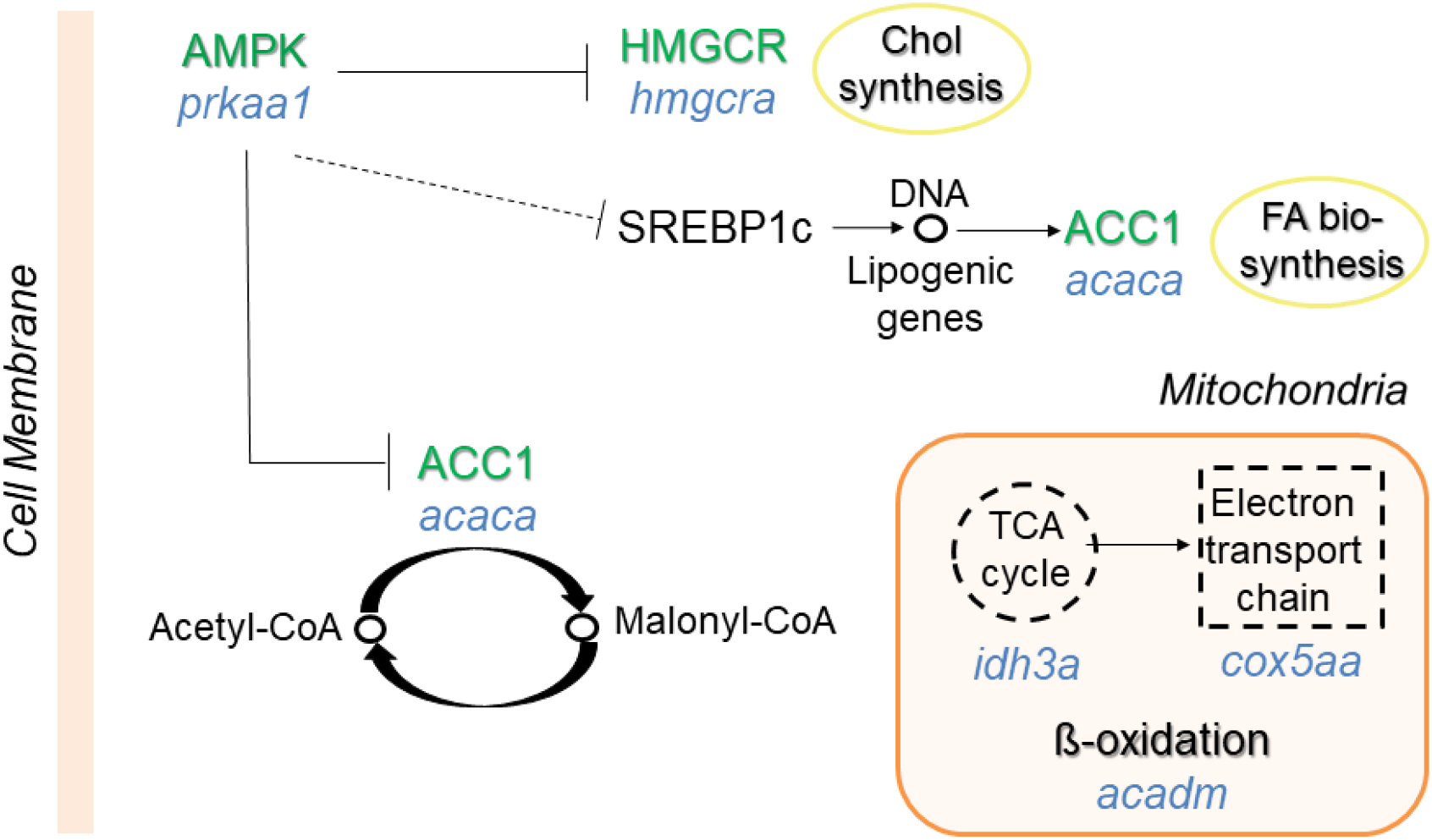
Summarized schematic representation of the main signaling pathways of the key selected genes involved in the putative MoA of MET and related to energy metabolism and lipid homeostasis. Green indicates key proteins in MET MoA, while blue indicates its encoding genes. AMPK – 5’ AMP-activated protein kinase; *prkaa1* – protein Kinase AMP-activated catalytic subunit alpha 1; HMGCR – 3-hydroxy-3-methylglutaryl-CoA reductase; *hmgcra* - 3-hydroxy-3-methylglutaryl-CoA reductase a; ACC1/*acaca* - acetyl-CoA carboxylase; SREBP1c - sterol regulatory element binding protein-1c; *idh3a* – isocitrate dehydrogenase (NAD(+)) 3 catalytic subunit alpha; *cox5aa* – cytochrome c oxidase subunit 5A; *acadm* – acyl-CoA dehydrogenase medium chain; Chol – cholesterol; FA – fatty acid; TCA – tricarboxylic acid cycle.

The qRT-PCR zebrafish primers for the genes *acaca, acadm, cox5aa, idh3a* and *hmgcra* were obtained in other published studies (Table 1). For the other genes, where the qRT-PCR primers were not available (the target genes – *prkaa1* and reference gene – *rpl13*), the zebrafish mRNA sequences were acquired from NCBI’s database (Table 1), and specific primers were designed for qRT-PCR using the Primer designing tool “Primer – BLAST” (NCBI) and Beacon Designer (Premier Biosoft International). To confirm the specificity of each primer, a PCR was performed in a Tgradient thermocycler (Biometra), and the subsequent products were run by electrophoresis in a 1.5% agarose gel. The resulting bands, with the expected size for each gene, were purified with GelPure (NzyTech), following the manufacturers’ protocol and sent for sequencing at GATC (Eurofins Genomics). The sequencing results were then uploaded and analyzed with the BLAST tool from NCBI in order to confirm the amplified sequence (Table 1).

**Table 1.**
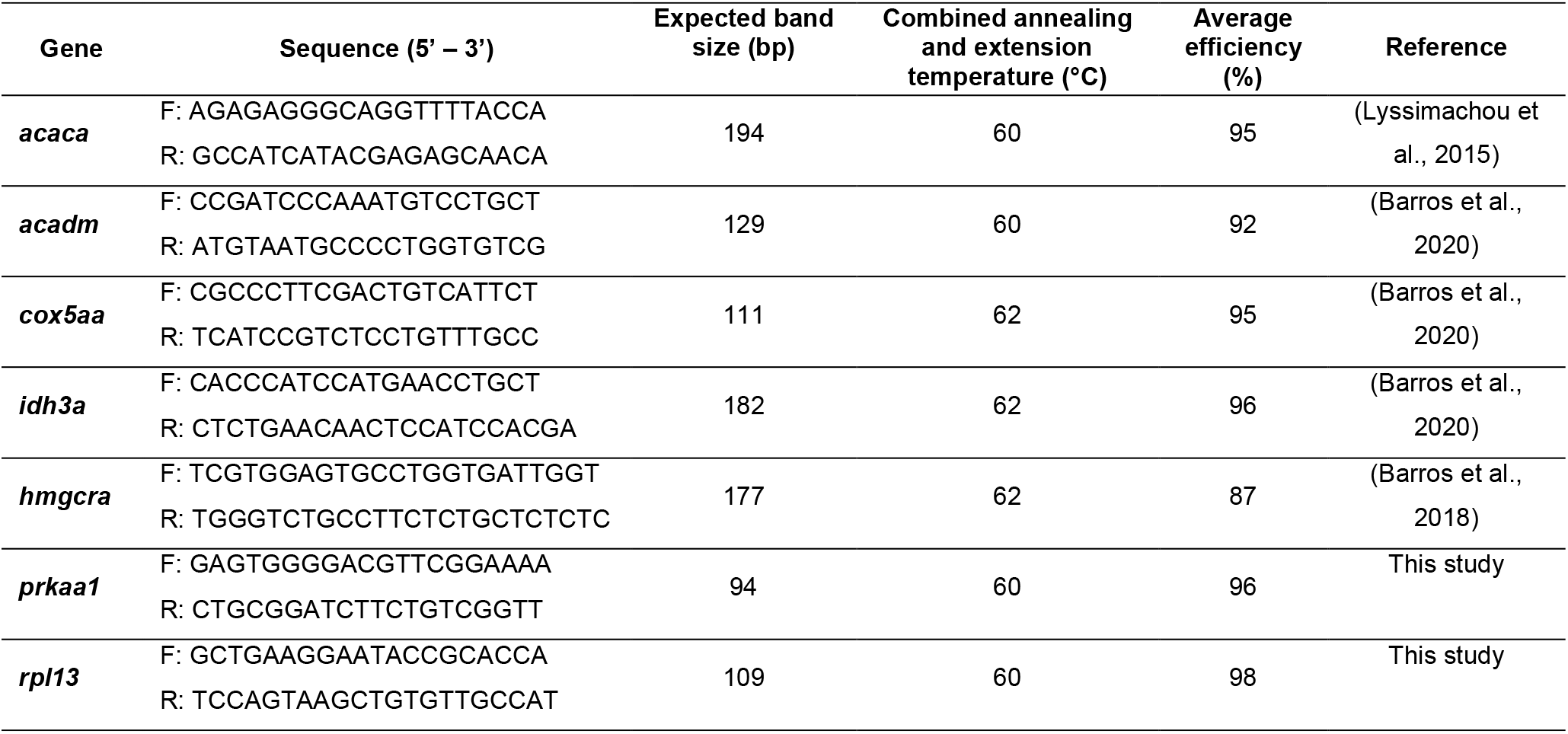
Primer sequences, forward (F) and reverse (R), and parameters used in the qRT-PCR for gene expression quantification in the 9-month-old zebrafish livers.

#### 2.8.4. cDNA synthesis and qRT-PCR analyses

Total cDNA was generated from 1 μg of total RNA previously extracted from 9 month-old adult livers (N=6-8/sex), using the iScript™ cDNA Synthesis Kit (Bio-Rad).

Gene expression profiles of *acaca, acadm, cox5aa, hmgcra, idh3a* and *prkaa1* were assessed by means of qRT-PCR. Ribosomal protein L13 (*rpl13*) was used as the reference gene since its expression levels did not show significant changes within the different treatments. Primers were already described by other authors or designed and tested for this study (Table 1). The cDNA of liver samples of each treatment were amplified in duplicate using a Mastercycler®ep realplex system (Eppendorf) in 96-well optical white plates, containing 5 µL of NZY qPCR Green Master Mix (2x) (nzytech), 0.4 µL of each primer at 400 nM (forward and reverse), 2.2 µL of water and 2 µL of cDNA at 100 nmol, in order to reach a final reaction volume of 10 µL. In every plate, a nontemplate control was included. The mRNA levels of selected genes were analyzed using a two-step qRT-PCR, which was performed as follows: 95°C of initial denaturation for 2 minutes, followed by 40 cycles of amplification with a denaturation at 95°C for 5 seconds and combined annealing and extension at 58 – 62°C, depending on the pair of primers previously validated, for 25 seconds. To confirm the specificity of the reactions, a melting curve (from 55 to 95°C) was generated in each run. PCR products were then analyzed by electrophoresis in 2% agarose gel to check the presence of single bands with expected size between 94 and 198 bp, depending on the pair of primers (Table 1), in order to confirm the specificity of the reaction. The PCR efficiency for the target and reference genes was determined by a standard curve, using six serial dilutions of cDNA pools of all samples (from 500 to 2.058 ng of cDNA in adult liver samples). The average PCR efficiencies obtained for target genes ranged from 87 to 98% (Table 1).

Relative change in transcription abundance of the genes of interest was normalized to *rpl13* and calculated using the 2^-ΔΔCt^ analysis method (Livak & Schmittgen, 2001). Control male and female expression levels (dechlorinated water treatment) were normalized to 1 and data were then expressed as fold changes to the control group.

### 2.9. Statistical analysis

Data obtained from this experimental study was checked for normality (Kolmogorov-Smirnov test) and homogeneity of variances (Levene’s test) followed by a one-way ANOVA. Post-hoc comparisons were carried out using Fisher’s least significant difference (LSD) test. Significant differences were set as p<0.05. All the statistical analyses were computed with Statistica 12 (Statsoft, USA).

## 3. Results

### 3.1. Analytical quantification of MET

The LC-MS/MS analysis revealed that no MET was detected in the control group. Average concentrations of 437.5 ± 62.6 and 342.5 ± 19.0 ng/L were detected in the 361 ng/L of MET treatment for the T1 and T2 sampling times, respectively. Due to technical problems, we were not able to quantify the T3 water sample from this treatment. Regarding the 2 166 ng/L treatment, actual concentrations quantified were 2 410.5 ± 18.1 ng/L (T1), 2 767.0 ± 368.4 ng/L (T2) and 3 610.6 ± 1 256.3 ng/L (T3) and in the 13 000 ng/L treatment, MET concentrations were 18 796.7 ± 186.0 ng/L (T1), 15 757.0 ± 1 182.4 ng/L (T2) and 8 718.0 ± 1 747 ng/L (T3). Overall, samples actual concentrations varied from their nominal ones by 28.8%, with a maximum variation of 66% in the T3 sampling time of the 2 166 ng/L treatment.

### 3.2. Mitochondrial complex I activity

The analysis on the activity of the mitochondrial complex I of the electron transport chain (COX I) revealed that MET significantly decreased this complex activity in males from all treatments (Figure 3). Additionally, females exposed to 2 166 and 13 000 ng/L of MET also showed decreased activity of COX I.

**Figure 3.**
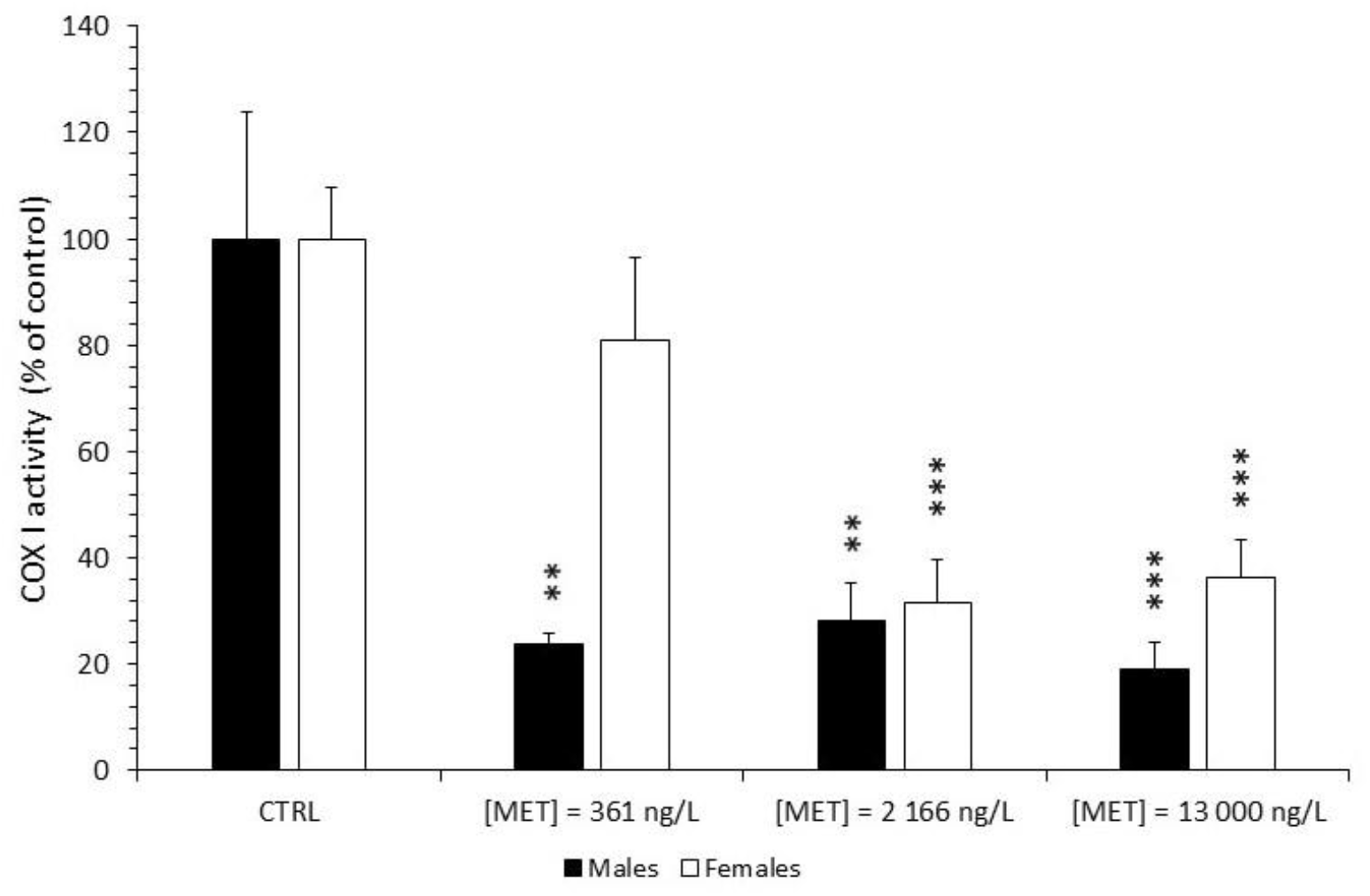
Mitochondrial complex I activity in adult *D. rerio* liver, expressed as percentage (%) of control, after 9 months of MET exposure. Error bars indicate standard errors; asterisks indicate significant differences from the control group (* - p < 0.05; ** - p < 0.01; *** - p < 0.001) (N = 5).

### 3.3. Cholesterol and Triglycerides quantification

Quantification of cholesterol (Figure 4.A) and triglycerides (Figure 4.B) in zebrafish males and females revealed significant differences from the control treatment, after 9 months of MET exposure. Cholesterol (Chol) concentration levels in the liver of the adult males, significantly decreased after exposure to 2166 ng/L and 13000 ng/L of MET, when compared with the control. Adult females also showed a similar pattern, being the cholesterol levels significantly lower for the intermediate concentration (2166 ng/L). Triglycerides’ (TGL) concentration in the adult male livers were significantly lower after exposure to the intermediate MET concentration (2 166 ng/L), in comparison to the control, whereas no effects were recorded for the lower (361 ng/L) and highest (13 000 ng/L) MET concentrations. This response appeared to be non-monotonic, showing an inverted U-shape. Females’ TGL remained unchanged.

**Figure 4.**
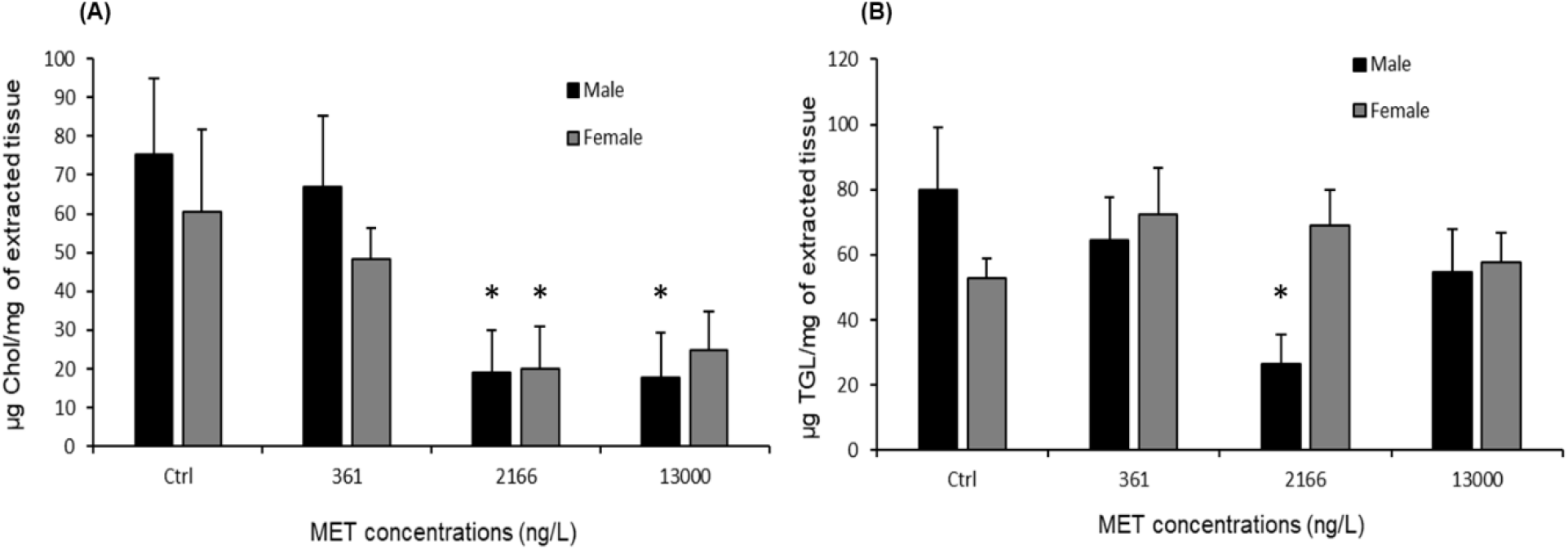
Effects of MET on *D. rerio* liver cholesterol (A) and triglycerides (B) levels, expressed as µg per mg of extracted tissue, after 9 months of exposure. Error bars indicate standard errors; asterisks (*) indicate significant differences from the control group (p<0.05) (N = 6-8).

### 3.4. RNA-seq

The RNA-seq analysis on zebrafish liver exposed to 13 000 ng/L of MET identified a total of 3 525 and 503 differentially expressed genes in males and females, respectively (Figure 5.A), with a false discovery rate (FRD) p-value (padj) < 0.005 and a log_2_(FoldChange) ≥ 1.5. Of these, 464 were upregulated and 3 060 downregulated in males (Figure 5.A), while 171 and 332 were upregulated and downregulated, respectively, in females exposed to 13 000 ng/L of MET (Figure 5.B). Additionally, the Venn diagram (Figure 5.C) revealed 188 common differentially expressed genes between males and females.

**Figure 5.**
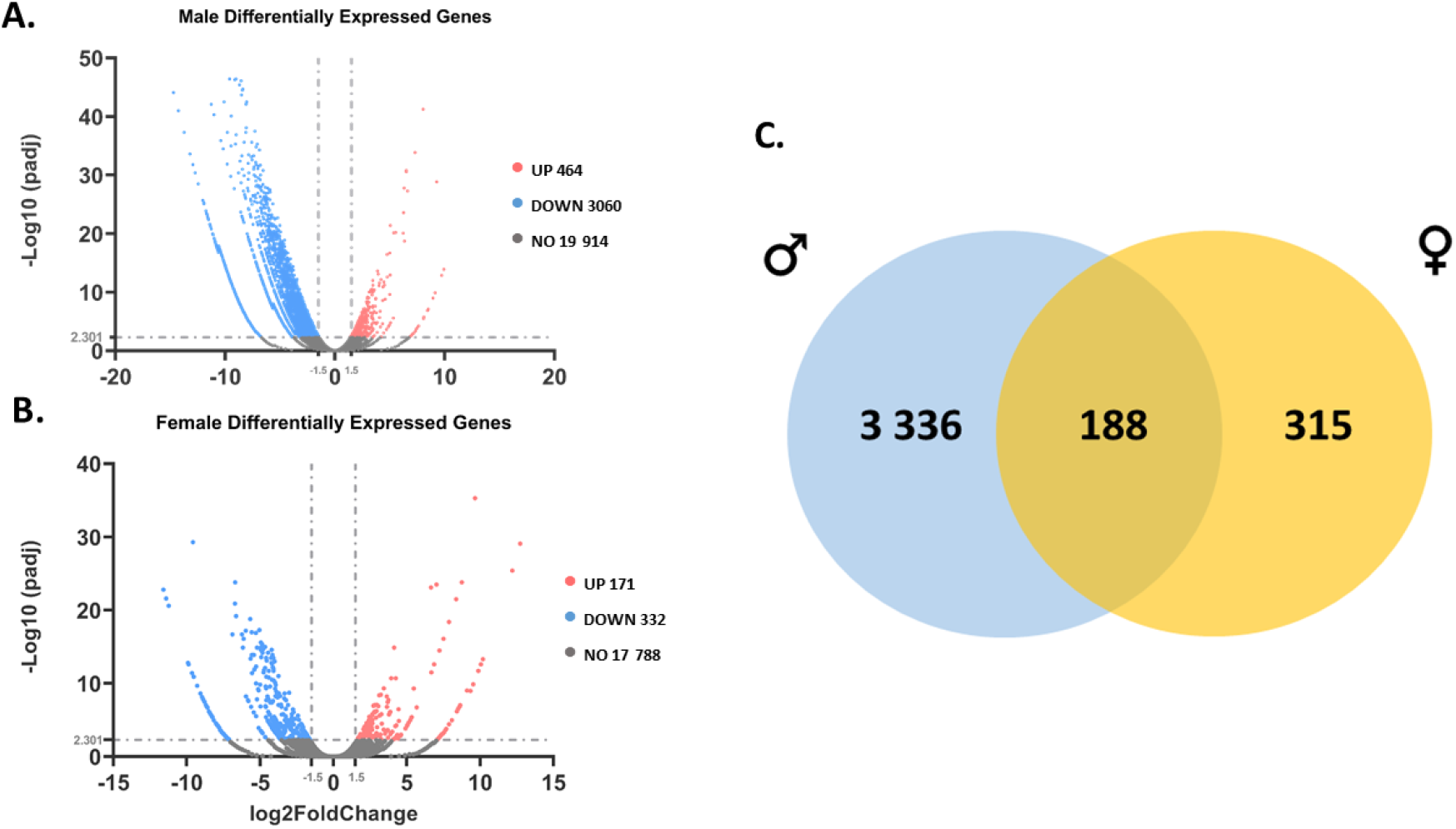
Transcriptomic analysis. Volcano plot on the differentially expressed genes in males (A) and females (B) hepatic tissue pools with 5 livers of *D. rerio* after a 9-month exposure to 13 000 ng/L of MET (N=1). padj ≤ 0.005 and a |log_2_(FoldChange)| ≥ 1.5. (C) – Venn diagram of the overlapping differentially expressed genes in males and females exposed to 13 000 ng/L of MET. UP – upregulation, DOWN – down regulation; NO – genes not differentially expressed; ♂ - males; ♀ - females.

A closer analysis on the genes involved in lipid and energy metabolisms was performed, which revealed 78 differentially expressed genes in exposed males, with most of them being downregulated, i.e. 55, and 23 upregulated (Figure 6.A). The analysis also revealed 44 altered genes in the liver of exposed females, 23 being upregulated and 21 downregulated (Figure 6.B). Moreover, only 8 genes had their mRNA levels altered in both sexes, i.e. *cyp11a1, cyp51, fabp2, ndufab1a, nme2b.2, scd, sqlea*, and *tpi1a*.

**Figure 6.**
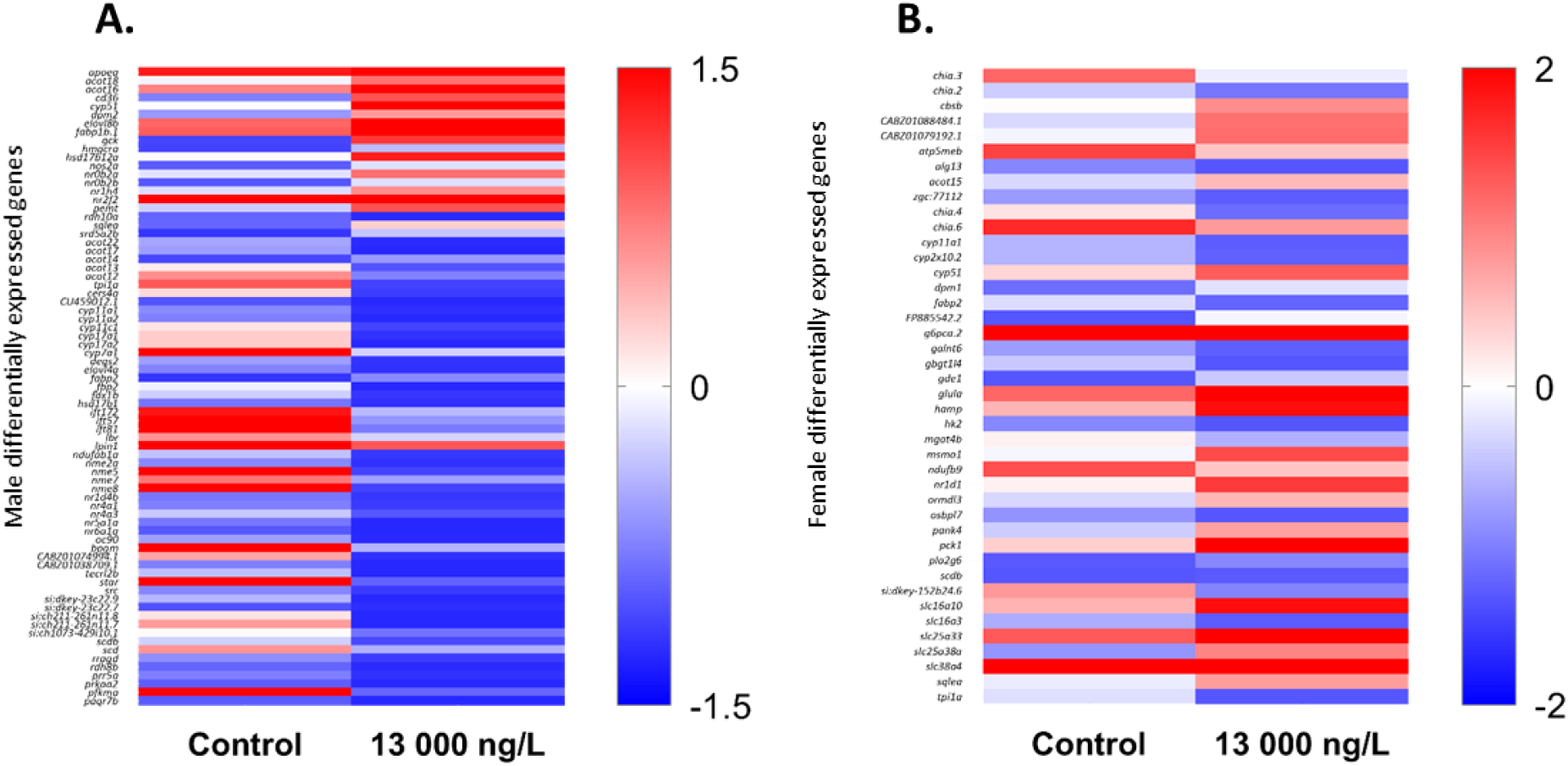
Hierarchical clustering heatmap depicting the expression patterns of gene transcription between control and 13 000 ng/L MET treatment. Data collected from male (A) and female (B) *D. rerio* liver pools (N=1), after a 9-month exposure to MET. Differentially expressed genes involved in the energy and lipid metabolisms are depicted. Data were clustered using the log_2_(FPKM+1) values. padj ≤ 0.005 and a |log_2_(FoldChange)| ≥ 1.5

### 3.5. qRT-PCR

The qRT-PCR analyses revealed that both male and female zebrafish livers (Figure 7 and 8) showed significant alterations from the control group on their gene’s expression levels after 9 months of exposure to MET, with females exhibiting more changes than males. In females (Figure 7), the *acaca* gene was significantly up regulated by 3.05 fold from the control group after exposure to 2 166 ng/L of MET. The transcription levels of *cox5aa* showed an up regulation of 1.45 fold for 364 ng/L of MET exposure. *idh3a* was also up regulated by 1.81 and 2.53 fold from the control treatment after exposure to 364 ng/L and 2 166 ng/L of MET, respectively. *hmgcra* expression was found to be highly up regulated in females, with a 4.73 and 13.45-fold increase from the control group after exposure to 364 ng/L and 2 166 ng/L of MET. On the other hand, a 5.88-fold downregulation of *hmgcra* was observed in males exposed to 2 166 ng/L of MET (Figure 8). In both male and female livers, the mRNA levels of all significantly altered genes followed a non-monotonic dose-response curve (NMDRC), with a U-shaped curve in males, and inverted U-shaped curve in females. Neither females nor males exhibited significant differences in *acadm* and *prkaa1* mRNA expression levels.

**Figure 7.**
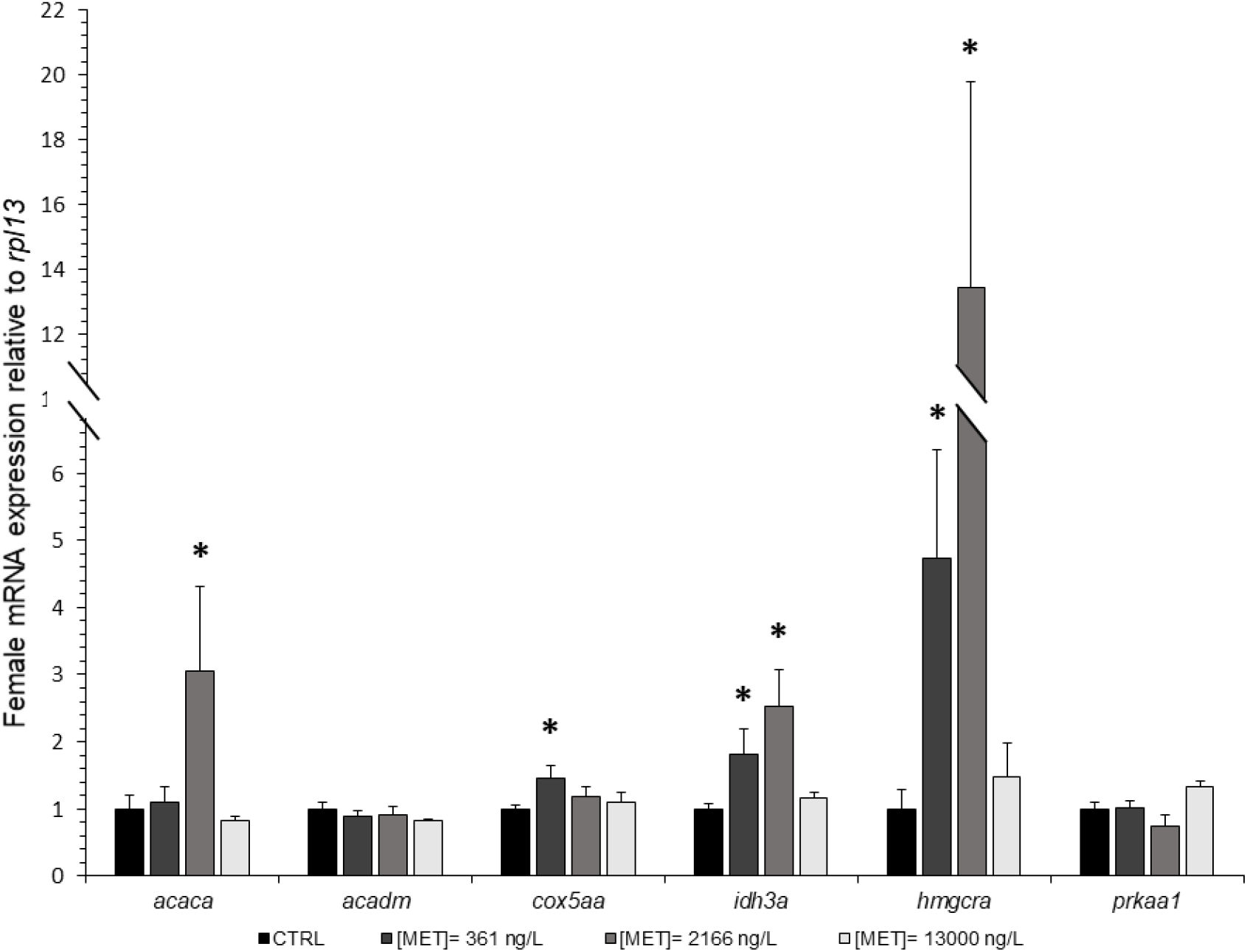
Females’ relative gene expression of acaca, acadm, cox5aa, idh3a, hmgcra and prkaa1, in the liver of adult D. rerio, after 9 months of MET exposure. Error bars indicate standard errors; asterisks (*) indicate significant differences from the control group (p<0.05) (N =6-8).

**Figure 8.**
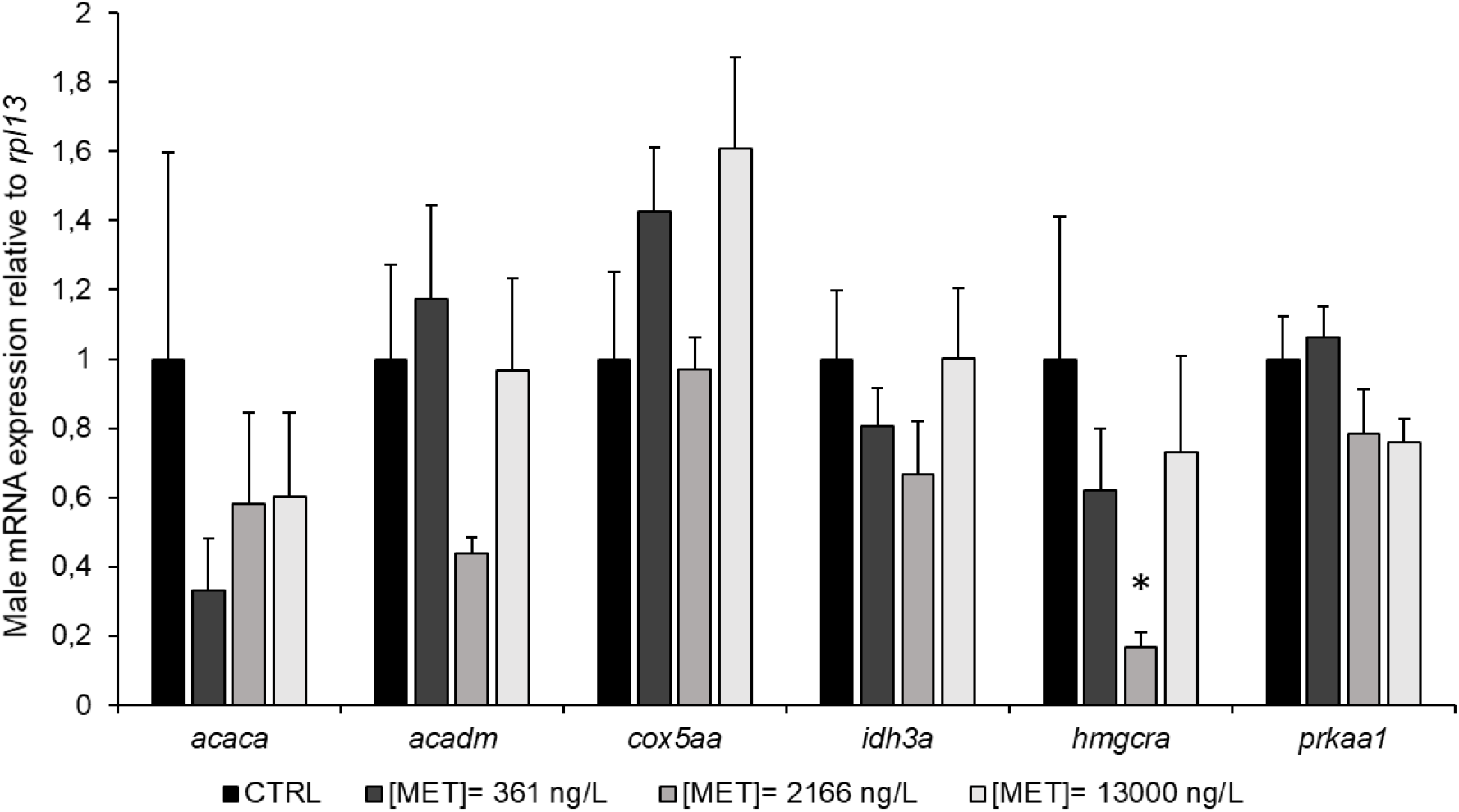
Males’ relative gene expression of *acaca, acadm, cox5aa, idh3a, hmgcra* and *prkaa1*, in the liver of adult *D. rerio*, after 9 months of MET exposure. Error bars indicate standard errors; asterisks (*) indicate significant differences from the control group (p<0.05) (N =6-8).

## 4. Discussion

MET is a pharmacologically active compound (PhAC); therefore, it is meant to produce biological responses at low concentrations potentially leaving non-target organisms at risk, particularly those that share conserved signaling pathways with mammalian, as the case of *D. rerio* (Arnold et al., 2014). Regardless of the growing number of studies focusing on the acute toxicity of MET in different aquatic organisms, there is a major knowledge gap concerning the generational effects of this contaminant of emerging concern in non-target organisms (Ussery et al., 2018). In order to address this knowledge gap, the present study aimed to investigate biological responses at biochemical and molecular levels, in the model organism zebrafish, after a generational exposure of 9 months to environmentally relevant concentrations of MET (361, 2 166 and 13 000 ng/L). This work integrated the determination of COX I activity, Chol and TGL quantification, transcriptomic analysis and the expression levels of key genes related to the putative MoA of MET using qRT-PCR, in livers of adult zebrafish to better understand the impact of MET in the lipid and energy metabolisms.

The results of this study showed that the 9-month chronic exposure to environmentally relevant concentrations of MET, was able to induce a series of disruptive effects on both biochemical and molecular endpoints. Furthermore, our parallel study (Barros et al., *in prep*) that addressed the effects MET in zebrafish morphometric and reproductive endpoints, reported increased hepatosomatic index - HSI in females (Figure 9). Results also revealed sex-dependent differences, which can suggest that females and males may not respond similarly to MET (Figure 9).

**Figure 9.**
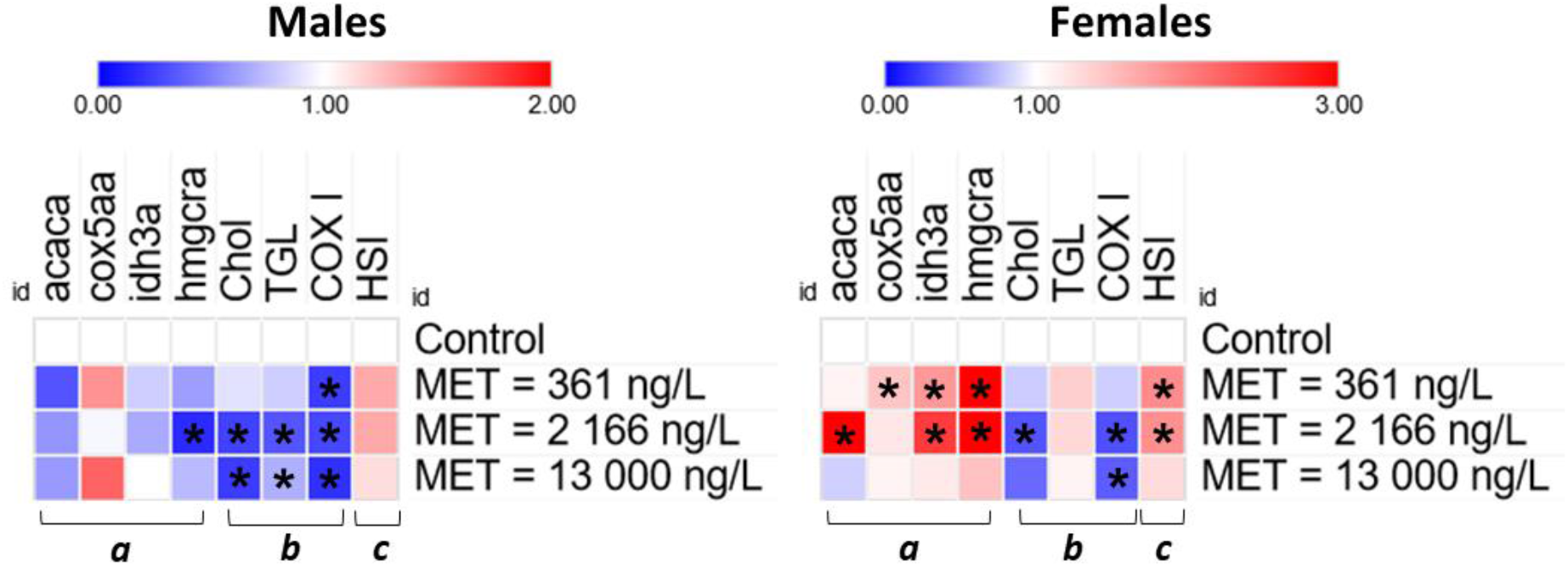
Summary of the effects of 9 months’ MET exposure on *D. rerio*. Asterisks (*) indicate significant differences from the control (p < 0.05). *hmgcra* – 3-hydroxy-3-methylglutaryl-CoA reductase a; *acaca* – acetyl-CoA Carboxylase Alpha; *cox5aa* – cytochrome c oxidase subunit 5Aa; Chol – cholesterol; TGL – Triglycerides; COX I – mitochondrial complex I; HIS – hepatosomatic index; a – gene expression; b – biochemical data; c – hepatosomatic index (HIS) from Barros et al. (*in prep*).

The biochemical analysis performed on the liver of exposed zebrafish revealed that MET was able to significantly decrease COX I activity at concentrations as low as 361 ng/L and 2 166 ng/L in both males and females, respectively (Figures 3 and 9). Additionally, the results of Chol and TGL quantification in males revealed that MET was effective in significantly lowering both parameters, while in females, MET was only effective in lowering Chol levels (Figures 4 and 9). These results are in agreement to the putative MoA reported in humans suffering from T2DM, where MET is described to inhibit COX I activity, as well as the synthesis of both Chol and TGL (Adak et al., 2018; Gong et al., 2012). Nevertheless, these effects can be harmful to non-target healthy organisms, where such a severe reduction of Chol and TGL may lead to adverse outcomes, as these molecules are essential for the organism homeostasis and energy reserves. Similar results on Chol and TGL levels were also found in a study with another fish species, *Nothobranchius guentheri* (Wei et al., 2020). However, the above-mentioned study used a MET concentration higher than those tested in our study (2 mg/g food). Interestingly, the present study showed that MET was more effective in lowering TGL levels in male zebrafish than in females. Also, COX I activity was affected at lower concentrations in exposed males.

In order to complement these biochemical results, two molecular analyses were performed: (1) RNA-seq and (2) qRT-PCR. The results of the transcriptome analysis (RNA-seq) in hepatic tissues are in line with the results here observed in biochemical endpoints. The analysis revealed several differentially expressed genes involved in lipid homeostasis and energy metabolism, in both sexes (Figure 6.A and B). However, these results should be carefully considered, since they are a result of an exploratory analysis of the animal’s hepatic transcriptome. Taking into account the overall results from the RNA-seq and biochemical analysis, several key genes, directly or indirectly involved in the putative MoA of MET, were selected for qRT-PCR analysis. At the end of 9-month exposure to MET, zebrafish presented alterations in the mRNA levels of *acaca, idh3a, cox5aa* and *hmgcra* genes. Interestingly, taking together with the results of biochemical parameters, it is possible to conclude that the intermediate concentration of MET (2 166 ng/L) was responsible for most of the significant changes in both analyses.

Regarding the existing literature, since MET is expected to decrease cellular ATP levels; and glycolysis is an important source of energy as the primary step of cellular respiration, it is expected that glycolysis emerges as a compensatory pathway (De Souza Silva et al., 2010; El-Mir et al., 2000; Foretz et al., 2010). mRNA transcription levels of *idh3a* and *cox5aa* reveal an upregulation in female livers, which indicate alterations in the tricarboxylic acid cycle and the electron transport chain, respectively, which may be a consequence of a stimulation of glycolysis.

Since COX I activity was found to be decreased in the present study, it is expected that ATP and AMP levels will change and consequently activate AMPK. Such activation leads to direct phosphorylation and inhibition of the limiting protein of the cholesterol pathway – HMGCR (Figure 1) (Gong et al., 2012). Even though *prkaa1* transcriptional levels were not significantly altered, transcription of *hmgcra* and Chol levels indeed were.

We hypothesized that the mRNA expression of *hmgcra* would be increased, as a negative feedback response to lower levels of cholesterol. In fact, females showed an upregulation of *hmgcra*, accompanied by a decrease in liver Chol levels. However, although male Chol levels decreased as anticipated, a downregulation of *hmgcra* was observed at the intermediate concentration of MET (2 166 ng/L). The expression profiles of *hmgcra* obtained for males in the present study may suggest that the low MET concentrations used in this experimental work affects zebrafish transcription levels of *hmgcra* in a time-specific manner, i.e., fluctuations in gene expression over time (Barros et al., 2018). Several studies have already addressed and described this time-dependent gene expression response, in a variety of aquatic organisms after exposure to different compounds (Cunha et al., 2017; Hamadeh et al., 2002; Kim et al., 2015). Therefore, the absence of studies on the temporal gene expression of *hmgcra* and other genes involved in the MoA of MET (i.e *prkaa1*), emphasizes the importance of using multiple time-points in gene transcription studies with this pharmaceutical.

No differences were observed in the results of mRNA expression levels of *acaca* (enables ACC activity) and *acadm* (catalyzes fatty acids ß-oxidation) in adult males, contrary to what was expected, as TGL levels were significantly lower. However, *acaca* mRNA transcription levels were significantly upregulated in females, even though a downregulation of this gene was hypothesized along with a decrease in TGL content, also not verified (Fullerton et al., 2013; Gong et al., 2012; Zhou et al., 2001). Some of the contrasting responses between mammalian animal models and zebrafish could be related with the well-established phenotypic differences between these two groups in regard to lipid metabolism and energy reserves, as fish tend to be more dependent on their lipid reserves than mammals (Polakof et al., 2012). Additionally, the differences between sexes here observed may be related to the fact that females have much larger fat reserves and higher lipid content than males, since females are responsible for egg production. This can be translated in higher energy demands and probably different gene expression profiles, between sexes, over time (Barros et al., 2018; Li et al., 2019). Therefore, it is possible that the gene expression profiles here observed are a result of both, gene expression fluctuations over time, as well as natural differences in the lipid and energy metabolisms between males and females. Overall, our qRT-PCR analysis reported that at the molecular level females were more sensitive to MET than males, mainly at the lower and/or intermediate concentration (361 and 2 166 ng/L), which integrates well with the significant increase of the HSI in females at the same concentrations, observed in our parallel study (Figure 9), indicative of metabolic disturbances (Lyssimachou et al., 2015).

The overall effects reported in our study may indicate an impairment of metabolic homeostasis in animals chronically exposed to MET. For instance, cholesterol and triglycerides are extremely important for the well-being of many aquatic organisms, including the zebrafish, due to their role in the maintenance of various biological processes, such as growth and reproduction (Barros et al., 2018). Disruption of energy and lipid metabolism, as observed in this study by the decrease in COX I, Chol and TGL levels and changes in the expression of genes involved in those processes, may imply harmful consequences for the organism and its population living in aquatic ecosystems. This is further corroborated by the significant increase in the females HSI observed in our parallel study, indicative of development of hepatomegaly (Figure 9; Barros et al., *in prep*).

Together with recent published studies, our data suggests that the current EQS and PNEC values (European Commission et al., 2022) underestimate the risk of MET in aquatic ecosystems as a concentration as low as 361 ng/L of MET was able to induce several detrimental effects in chronically exposed zebrafish. Therefore, there is an urgent need to review the EQS and PNEC values for MET, as concentrations tested in this study are all deemed safe by current EQS and PNEC values

Most of the present results of qRT-PCR and biochemical analysis demonstrated a non-monotonic dose-response curve (NMDRC) with an inverted U-shaped curve in females’ gene expression of *acaca, cox5aa, idh3a* and *hmgcra*, and a U-shaped for males’ gene expression of *hmgcra* and males TGL levels. For a long period of time it was assumed that contaminants would present a linear monotonic dose-response (Vandenberg et al., 2012). However, an increasing number of long-term studies, have documented the occurrence of NMDRC as a response to low concentrations of several compounds, mainly endocrine-disrupting chemicals (Andrade et al., 2006; Barros et al., 2020; Crépeaux et al., 2016; Kim et al., 2014). The mechanism(s) behind this non-monotonicity are not fully understood, however, the explanatory basis of NMDRCs is based on adaptation mechanisms, generally dependent on the exposure time and the range of concentrations tested. To date, there is no record of this type of response to MET, and thus, we raise the possibility that this is the first study to demonstrate NMDRCs in response to a full life-cycle exposure to environmentally relevant concentrations of MET.

## 5. Conclusion

The present study revealed several effects of MET on an aquatic model organism, still not reported in the literature for this pharmaceutical. The generational exposure to MET affected biochemical and molecular responses in the model organism *D. rerio*, including disruption of important mechanisms related with energy metabolism and lipid homeostasis along with changes at the individual level, such as HIS, potentially related with liver dysfunctions. Importantly, the data generated here is of great relevance to improve environmental risk assessment, given that aquatic organisms are chronically exposed to MET during multiple generations, at low concentrations. Moreover, the present data indicates that the current EQS and PNEC values for MET proposed in the draft of the 4^th^ watch list should be revised, as in this study the concentrations as low as 361 ng/L produced important detrimental effects in zebrafish metabolism and indicate potential liver dysfunctions. Further studies should be performed on the long-term effects of MET exposure at environmentally relevant concentrations in other taxa, since most of the ecotoxicological studies are limited to aquatic vertebrates.

## 6. Acknowledgements

This study was developed under the project Nor-Water – Pollutants of emerging concern in watersheds from Galicia-northern Portugal: new tools for risk management [Reference: 0725_NOR_WATER_1_P], financed by Programa de Cooperação Interreg Portugal/Espanha, (POCTEP) 2014–2020. The study was also supported by the National Funds through Portuguese Foundation for Science and Technology (FCT) under the projects [UIDB/04423/2020, UIDP/04423/2020) and UIDB/04033/2020]. S. Barros, M. Pinheiro, H. Morais and N. Alves acknowledge FCT for their Ph.D. grants PD/BD/143090/2018; SFRH/BD/147834/2019; SFRH/BD/139762/2018 and DFA/BD/6218/202, respectively. Financial support by Xunta de Galicia (ED431C 2021/06) and the Spanish Agencia Estatal de Investigación - MCIN/AEI/ 10.13039/501100011033 (PID2020–117686RB-C32) is also gratefully acknowledged.

